# Genetic basis of a seasonal life-history polyphenism

**DOI:** 10.1101/2022.09.30.510344

**Authors:** Serena A Caplins

## Abstract

Seasonal polyphenisms are common across the animal and plant kingdom yet we understand the explicit interactions between genetics and environment for only a few taxa. Are the genomic regions and their variants associated with the trait the same or different across environments? Is the response to selection shared or different across different “background” selection environments? Offspring type in the sacoglossan sea slug *Alderia willowi* is a seasonally modulated interaction between genotype and phenotype that results in offspring of wildly different developmental trajectories. In a genome-wide association test I found 41 SNPs associated with offspring type. In an evolve and resequence experiment I found thousands of loci changed in frequency following selection. These loci were partially shared (37%) between low and high salinity. Of the 41 candidate SNPs identified in the GWAS only seven also showed significant allele frequency change across replicates in the selection experiments with four in high salinity, two in low and one in both. This reveals a broad pattern of allele frequency change that is largely unique to the environment in which selection for the same phenotype occurs. The results presented in this paper showcase the ability of phenotypic plasticity to move the phenotype independent of the genotype and thus maintain the polyphenism that is so striking in this species.

## Introduction

Polyphenisms are a form of adaptive phenotypic plasticity where the same genotype can produce different discrete phenotypes when the environment varies. These include scale color in butterflies, wing-morph in crickets and aphids, and offspring type in sea slugs (Krug et al. 2012, Caplins 2021). Seasonal polyphenisms are a common form of polyphenism as seasonal conditions provide a reliable cue to which such discrete plastic response can evolve (Nijhout 2003). Along the Western North American coast, temperature and salinity vary seasonally in pronounced ways. Winter rainfall events can lower the salinity of estuarian and intertidal habitats to freshwater levels for several hours, and to brackish (a mixture of salt and freshwater) conditions for days (Hutton et al. 2015). Temperature also varies substantially, increasing in the summer months, the effects of which are felt most strongly during diurnal low tides. In this paper I explore the effects of salinity on the response to selection for offspring type in a sacoglossan sea slug which has shown seasonal local adaption to low salinity (Krug et al. 2021).

Marine invertebrate larvae are either maternally provisioned with the resources they need to proceed through development, or they must feed in the plankton to acquire these nutrients. These two modes of development, lecithotrophy and planktotrophy, respectively, are widespread across all major animal phyla, and are bimodally distributed with few intermediates (Strathmann 1978; Collin 2012, Collin and Moran 2017). The evolution of developmental mode has sparked decades of research to understand the selection pressures, genetic architecture, phylogenetic relationships, and ecological consequences of these pervasive traits (Strathmann 1978; Levin, Zhu, and Creed 1991; Marshall et al. 2012; Marshall and Burgess 2015; Zakas et al. 2018; A. F. Armstrong and Grosberg 2018). While species with a mode of development that is intermediate between lecithotrophy and planktotrophy are rare, there are a few exceptional systems in which are capable of producing both types of larvae either as a fixed polymorphism (Levin, Zhu, and Creed 1991; Zakas et al. 2018) or a seasonal polyphenism (Patrick J. Krug, Gordon, and Romero 2012; Collin 2012), or which produce facultatively feeding larvae (Armstrong and Lessios 2015). Poecilogonous species exhibit intraspecific variation for developmental mode and provide a clear way to explore the selection pressures and corresponding genetic and transcriptomic changes that drive the evolution of developmental mode (Levin, Zhu, and Creed 1991; Anne Frances Armstrong and Lessios 2015; Zakas et al. 2017). Poecilogonous species provide a view into the specific alternative developmental pathways that give rise to these modes (Knott and McHugh 2012; Zakas et al. 2018).

Understanding the genetic architecture (number and effect size of genes associated with a phenotype) of developmental mode will greatly add to our understanding of how it has evolved and how it will respond to future selective pressures. Evolve and resequence studies incorporate the power of artificial selection experiments and high throughput sequencing to distinguish the number and effect size of genes that respond to selection from those that change in frequency due to drift as the phenotype changes in response to artificially imposed selection (Kawecki et al. 2012; Long et al. 2015; Schlötterer et al. 2016). Evolve and resequence studies can also be used to address the repeatability of evolution through replication of selection lines (Kawecki et al. 2012), and to test the impacts of environmental factors as selective agents or as cues for phenotypically plastic traits. Using an evolve and resequence approach has been particularly useful in identifying the genetic basis of many polygenic traits, including body size (Turner et al. 2011) courtship song (Turner and Miller 2012), egg size (Jha et al. 2015), aging (Remolina, Chang, and Leips 2012), response to climate change (Brennan et al. 2020), and adaptation to plant host in seed beetles (Gompert and Messina 2016).

*Alderia willowi* is a sacoglossan sea slug that exhibits a seasonal polyphenism for developmental mode (Patrick J. Krug, Gordon, and Romero 2012). Field-collected individuals of *A. willowi* produce more lecithotrophic embryos during the summer than the winter (Patrick J. Krug, Gordon, and Romero 2012), a pattern that can be replicated in the lab under conditions mimicking mean summer and mean winter temperature and salinity regimes (Patrick J. Krug, Gordon, and Romero 2012). Previous studies show that developmental mode in *A. willowi* responds readily to selection for increased proportions of lecithotrophy but that the response to selection over one generation is reduced in low salinity due to maternal effects (Caplins, 2021). In *A. willowi*, egg size is correlated with developmental mode: egg diameters are bimodally distributed, and large eggs (X ± SD: 105 ± 5 µm) develop into lecithotrophic larvae that metamorphose into juvenile slugs in ∼5 days, whereas small eggs (X ± SD: 68 ± 4 µm) give rise to planktotrophic larvae that only become metamorphically competent after 21 days of feeding on planktonic algae <u>(P. J. Krug 1998)</u>. Both size classes of larvae can feed on phytoplankton, but the larger, lecithotrophically developing larvae do not need to feed to complete metamorphosis and occasionally develop into the juvenile stage while still encapsulated in their egg capsule, by-passing a swimming stage altogether <u>(P. J. Krug 2001; Botello and Krug 2006)</u>. *A. willowi* is a hermaphroditic, euryhaline species whose range extends along a seasonal temperature and salinity gradient in estuaries from San Francisco Bay, CA to Baja California, Mexico, where it is an obligate consumer of *Vaucheria* sp. algae. Freshwater input via winter rain results in up to a 50% reduction in salinity for northern populations lasting several months (∼December – March), as well as temporary hyperosmotic (∼2 ppt) conditions during low tide (Garchow and Krug 2010). Slugs exhibit a heritable tolerance to low salinity, with northern populations more tolerant of low salinity (2 ppt) than southern populations with respect to survival (Koch and Krug 2012). Salinity significantly influences egg-mass type, with slugs reared in high salinity more likely to lay lecithotrophic egg masses (Patrick J. Krug, Gordon, and Romero 2012, Caplins 2021). There is a significant interaction between egg-mass type, salinity, and family, suggesting egg-mass type is determined via a GXE interaction (Caplins 2021).

In this paper I first test for genomic associations with offspring type via a genome wide association study (GWAS) for individuals reared in the lab for one generation in high salinity. To determine whether the response to selection is shared across salinity environment I then select for increased proportions of lecithotrophy across replicate populations in low and high salinity to test the effect of salinity environment on the response to selection over 5 generations. I compare the SNPs from the GWAS with those from the selection experiment to determine if the same sites that were associated with the phenotype also change in frequency due to selection.

## Methods

### Experimental summary

To test the effect of the background selection environment on selection for increased lecithotrophy, I reared slugs in low and high salinity while selecting exclusively for lecithotrophy. To estimate the genetic architecture of egg-mass type, I used slugs from the first generation of the high salinity treatment to preform a genome wide association study for egg-mass type (lecithotrophic or planktotrophic producing). Sequencing slugs for the GWAS also provided me with pre-selection samples to compare with generations post-selection to determine whether the alleles that change in frequency due to selection are shared across salinity environments. I also use the post-selection samples to test whether the alleles that change in frequency are the same as those found associated with the phenotype in the (pre-selection) GWAS.

A brief note of clarification: Confusion can arise from the nature of the egg-mass phenotype. As hermaphrodites, each slug is capable of producing an egg-mass and thus has an egg-mass phenotype. Each slug also has its own developmental history, i.e., whether that slug was lecithotrophic or planktotrophic in its own early development. For this paper I focus on the adult phenotype, which type of offspring an adult slug produces. For the selection experiment only lecithotrophically developing slugs survive each generation, thus all individuals share a common developmental history and it is only the adult phenotype of egg-mass type that varies for these experiments. Thus, unless explicitly stated, egg-mass type or offspring type in this paper refers to a character of the adult (what kind of offspring is produced).

### Sample collection and selection experiment

I collected approximately 1500 adult slugs from Long Beach, CA on 7 May 2018 [33.73 N, 118.203 W]. I transported the slugs to UC Davis where all the following experiments took place. Field-collected slugs were maintained in two 24 × 32 × 4 cm glass dishes with air stones at room temperature (20-22 °C), ambient sea water (32 ppt) and fed freshly collected *Vaucheria longicaulis* algae as they deposited egg masses. Over a four-day period I removed 400 lecithotrophic egg masses from the field collected slugs.

Juvenile slugs hatched from these 400 egg masses within three days. I haphazardly placed between 250-300 juvenile slugs into each of 6 selection lines. The selection lines include three replicate lecithotrophic lines in both low (16 ppt) and high (32 ppt) salinity. Lines were reared in 24 × 24 × 4 cm glass baking dishes, filled with 2 liters of water, fitted with 1 air stone, and covered in plastic wrap to minimize evaporation. Selection lines were housed in a walk-in environmental chamber maintained at 20°C. Within each selection line replicate, slugs were allowed to mate freely and deposit egg masses at will. Three times per week I fed the slugs freshly collected *V. longicaulis*, replaced 50% of the seawater in the dishes, and checked for egg-masses. When I observed an egg-mass, I placed it in a small glass finger bowl where I recorded the type of egg-mass (lecithotrophic or planktotrophic). The progeny from lecithotrophic egg-masses were used to start the next generation of each selection line, from which approximately 200 juvenile slugs were selected each generation from each line separately. Specifically, I held all lecithotrophic egg-masses in small glass finger bowls with an air stone, sea water respective to the salinity of each selection line (low or high), and fresh *V. longicaulis* to stimulate metamorphosis (Botello and Krug 2006), until the embryos developed into juveniles. When at the juvenile stage I haphazardly counted out 200 slugs for the next generation, respective to each line. In most cases 200 juveniles was near the maximum produced. At no point in time were the lecithotrophic larvae fed planktonic food, requiring that larval development be entirely lecithotrophic. A subset of adults was flash frozen (50-100 individuals) each generation. I calculated realized heritability on developmental mode for a given salinity using the breeder’s equation (R = h^2^S) modified for a threshold response using a probit transformation to translate the proportion of individuals expressing the trait of interest to a mean value for that trait (Walsh and Lynch 2018).

### Sample sequencing and processing

Samples were sequenced across three lanes (pre-selection: 2 lanes, post-selection: 1 lane). From the pre-selection treatments, I extracted DNA from 192 individuals using the pure gene protocol (Fisher Scientific). From the post-selection samples, I extracted DNA from 6-19 individuals per selection line (total: 96, mean ± SD: 12 ± 4.4). I prepared the DNA samples for sequencing using the restriction enzyme associated DNA markers (RAD-tags) BestRad protocol. I used the restriction enzyme *sbfl-hl* on 200 ng per ul DNA for each sample. For the pre-selection samples, I sonicated the pooled individually barcoded libraries to a mean fragment size of 500 base pairs as verified on a bioanalyzer DNA HS chip, using a using an NGS bioruptor for the pre-selection samples, and a Covaris e220 sonicator for the post-selection samples. The pooled libraries were sequenced on illumina HiSeq 4000 PE 150. I demultiplexed the sequences and removed PCR duplicates in stacks (v 2.41). I mapped the reads to the *A. willowi* reference genome (see below) with Bowtie 2 (v 2.3.4.1). I called genotype likelihood estimates in ANGSD (v 0.936) using the GATK model (McKenna et al. 2010) with a p-value threshold of 1e-6. To accounted for variation in genotype calls due to sequence lane (lane effect), I re-sequenced 14 samples from the pre-selection samples described above in the third illumina lane with the post-selection samples.

I estimated pairwise relatedness within each treatment and generation (f1: high salinity, f5: low and high). I used ANGSD to generate genotype likelihoods with a minimum SNP pvalue of 1e-6, and a minimum minor allele frequency of 0.05. I then used NGSRelate (v2, Hanghoj et al. 2019) to calculate pairwise relatedness (Hedrick and Lacy, 2015). I included the 14 resequenced samples from the f1 high salinity treatment which were resequenced to ensure consistent SNP calling between sequencing libraries and lanes. These samples should have a relatedness of 1 to themselves and provide a positive control for the relatedness estimates.

### GWAS

To test for genetic variants associated with egg-mass type (planktotrophic or lecithotrophic), I phenotyped high salinity F1 individuals from the selection experiment by placing individuals in 12-well culture plates, one individual per well and of these I sequenced 192 samples (plus 14 re-sequenced individuals) as described above. I used the genotype likelihood estimates to run a principal component analysis in NGS tools (Fumagalli et al. 2013). I filtered the SNPs in ANGSD (v 0.939) by only retaining those with a p-value 1^-6, or less, and imputed genotype likelihoods in beagle (v3). I then ran a genome wide association test in ANGSD using imputed genotype likelihood scores against whether a slug had laid a lecithotrophic egg mass (phenotype = 1) or a planktotrophic egg mass (phenotype = 0). To account for the effect of relatedness, I took the first eigen vector of the pairwise relatedness estimates (described above) and included it as a covariate in the GWAS. I also included sequencing lane as a binary coded covariate. To test the validity of the GWAS against the real phenotype, I re-ran the genome wide association test using the above-mentioned covariates on ten iterations of a randomly assigned phenotype. I compared the values for the likelihood ratio test statistic from these ten random iterations against the real phenotype.

The likelihood ratio statistic is chi-square distributed with one degree of freedom and is derived from a linear regression which provides a likelihood ratio score for each SNP (Skotte et al. 2012). I adjusted p-values for multiple testing using q-values (FDR = 0.1), Benjamin Hochberg (alpha = 0.1), and bonferroni (alpha = 0.1).

### Cochran-Mantel-Haenszel test

To test the degree to which the change in allele frequency following selection for increased lecithotrophy was shared between the salinity treatments I performed a Cochran-Mantel-Haenszel (CMH) test on allele count. I also used the results of the CMH test to determine whether the top loci that changed in allele frequency are the same as the candidates identified in the GWAS. I used several R packages (“poolSeq”, “BiocManager”, “qvalue”, “ggplot2”, and “ggvenn”). I calculated allele frequencies for each replicate line from pre and post-sequenced samples in ANGSD (v 0.938). Allele frequencies were estimated assuming known major and minor allele and taking the uncertainty of the minor allele into account via genotype likelihoods (-doMaf 3) using the GATK estimator (-GL 2). I used these allele frequency estimates multiplied by coverage (number of individuals at a given locus) to calculate allele count. I performed the CMH test for replicate samples in low and high salinity separately to determine the degree of similarity in the response to selection across the two salinity environments. I corrected the resultant p-values from these tests for multiple testing using an FDR of 10%.

### Reference genome assembly

A *de novo* reference genome was constructed for *A. willowi* using 10X genomics from a slug collected from Mill Valley, CA [37°52’55”N 122°31’03”W] on 12 September 2017. This individual was flash froze in liquid nitrogen 4 days after collection to limit the amount of contamination from food remaining in the digestive tract. I extracted DNA from a single whole individual using the salting out protocol (10X Genomics), which I modified for small amounts of tissue (github: SerenaCaplins). I sent the resulting high molecular weight DNA sample to the UC Davis Genome Center for library preparation and paired-end 150 base pair sequencing on an Illumina HiSeq 4000 (Davis, CA). I assembled the sequence data in supernova (v 2.0.1) using 50% of the sequenced reads to obtain the recommended coverage for which supernova is optimized (10X Genomics support staff personal communication). I generated an output fasta file using the pseudohap style parameter in supernova, which generates a single record per scaffold (10x Genomics). I evaluated assembly completeness in BUSCO (v 1.22) using the metazoan database.

## Results

### Selection experiment

The parental generation laid 36% (80/217) lecithotrophic egg masses within the first 24 hours of being in the lab. I started the F1 generation with an average of 250 slugs across two treatments (salinity: low versus high) replicated 3 times. The average number of slugs per replicate in generations F2-F5 was 218 (Range: 160-263) The mean number of lecithotrophic egg masses each generation across all replicates ranged from 20 to 197 with a mean of 68 ± 45 (SD). The proportion of lecithotrophy per replicate increased steadily for most replicate selection lines, on average increasing from 32% to 64% in low salinity and from 43% to 56% in high salinity. Based on a probit transformation to translate the proportion of individuals expressing the trait of interest to a mean value for that trait (Walsh and Lynch 2018), estimates of realized heritability for the proportion of lecithotrophy were 0.16 in high salinity and 0.31 in low salinity. In the first two generations, more lecithotrophic egg-masses were produced in high salinity than low salinity. This difference was no longer present by the third generation (Figure 2).

**Figure 1.**
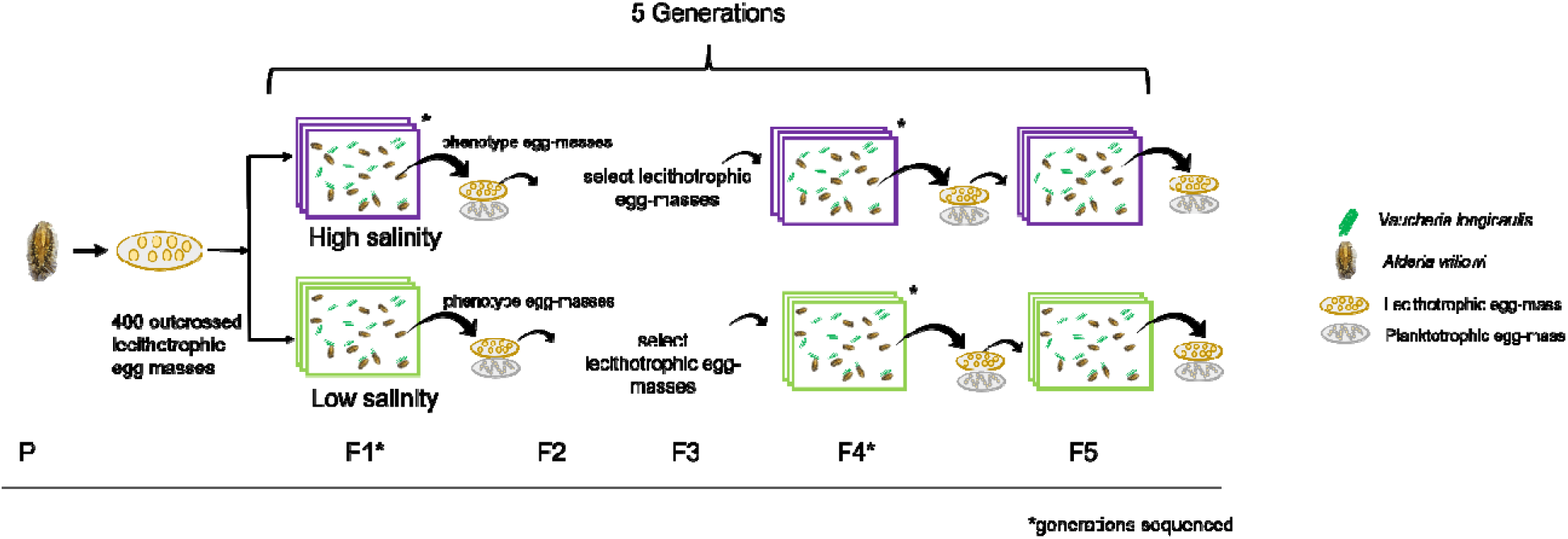
Experimental schematic. Adult slugs were collected from Long Beach, CA. The lecithotrophically developing larvae which emerged from their egg-masses (N = 400) were evenly divided into the low and high salinity treatments, with ∼200 slugs in each treatment, replicated three times. Each generation the proportion of lecithotrophic egg-mass type was determined and lecithotrophic egg-masses selected for the next generation. Pre-selection samples were composed of slugs from the F1 generation in high salinity and were used for a GWAS and baseline allele-frequency. Post-selection samples were from the F4 generation in low and high salinity and were sequenced to examine the genomic response to selection.

**Figure 2.**
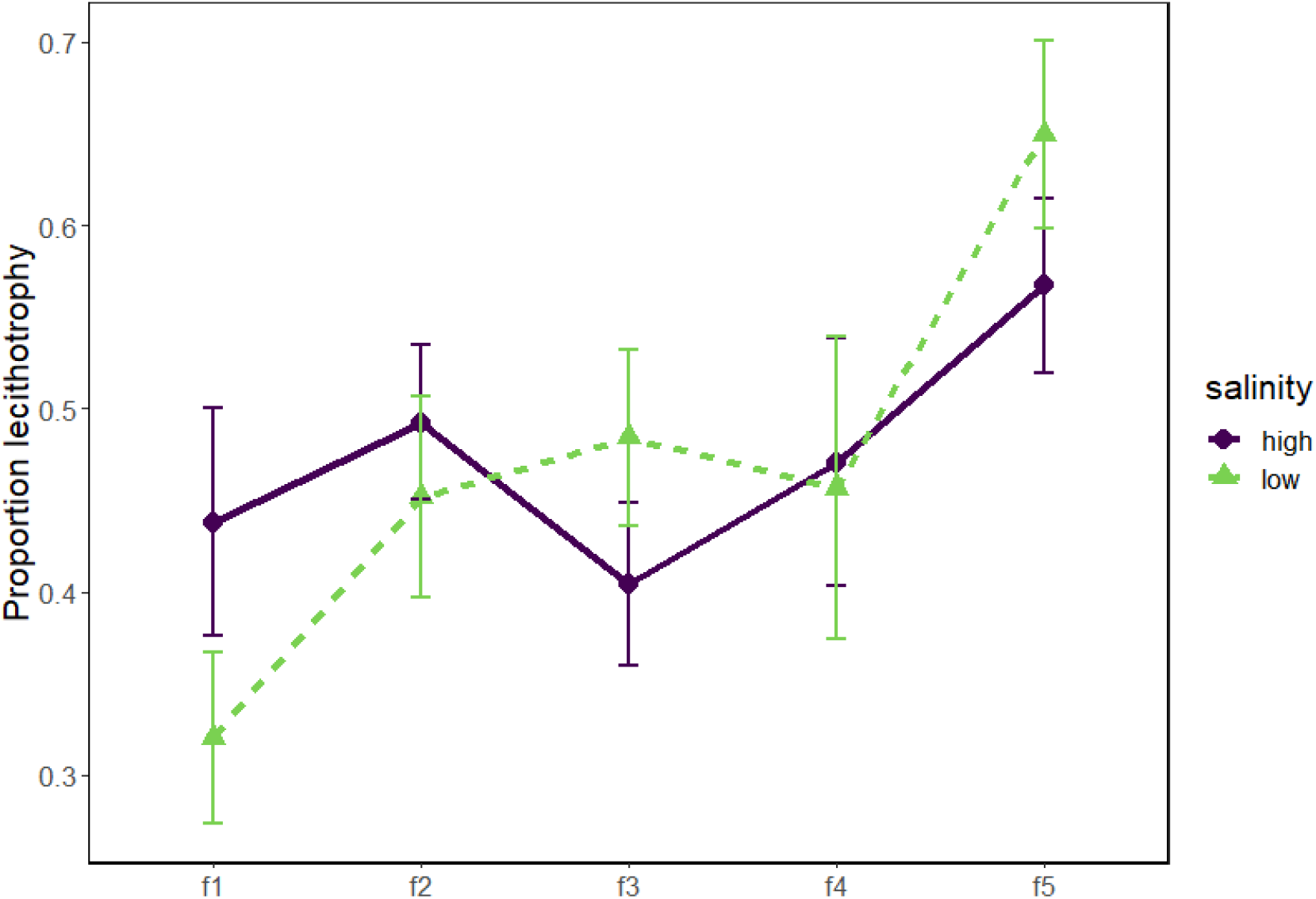
Phenotypic response to selection for lecithotrophy for 5 generations in low (16 ppt) and high (32 ppt) salinity.

### Sample sequencing and processing

I obtained an average of 300 thousand reads per sample after filtering (removing PCR duplicates and low-quality mapping reads), resulting in an effective per-sample coverage of 130x (mean, sd = 109x, min = 1x, max = 639x) as estimated in gstacks (v2.4). Nine individuals were excluded from the pre-selection samples, and 18 from post-selection due to a low number of paired reads (< 20k) and low effective coverage. For the pairwise relatedness estimates I excluded estimates of relatedness (rab, Hedrick and Lacy 2015) that used less than 40k sites. Mean pairwise relatedness for pre-and post-selection samples was centered around zero, though it did increase following selection, with preselection = 0.0066 (sd = 0.0066), post-selection low salinity = 0.033 (sd = 0.093), post-selection high salinity = 0.018 (sd = 0.065). A t-test between pre-selection and low and high salinity found no significant difference between pre and high (t = -1.95, df=127.7, p-value = 0.053), but a significant difference with low salinity having higher relatedness (t = -4.19, df = 222, p-value = 3.885 e -05). The 14 individuals from pre-selection that were re-sequenced with the post-selection samples had a relatedness value between 0.99 and 1. Excluding the resequenced samples, there were three pairs of individuals in the pre-selection samples with a relatedness greater than 0.5, and two pairs in the low salinity post-selection samples. One individual from each pair was removed from subsequent analyses. Filtering by read count, and relatedness left 187 individuals’ pre-selection and 64 post-selection. In a PCA of the pre-selection samples 19% of the variation explained in the first two PCs, there was no grouping of individuals according to whether they laid lecithotrophic or planktotrophic egg-masses (Figure S1).

### Common Garden GWAS

Of the 186 pre-selection slugs, 85 laid lecithotrophic egg masses and 101 laid planktotrophic. I calculated genotype likelihood scores for 1,397,808 sites. The genome wide association test returned LRT values for 67,871 SNPs post-filtering which are distributed across 2,848 scaffolds. There were 41 SNPs with an LRT value greater than 15 (max LRT = 28), 27 of these were significant by q-value, 13 by BH, and 4 by bonferroni correction for multiple testing (Figure 3). In the GWAS randomizations I found an average of

**Figure 3.**
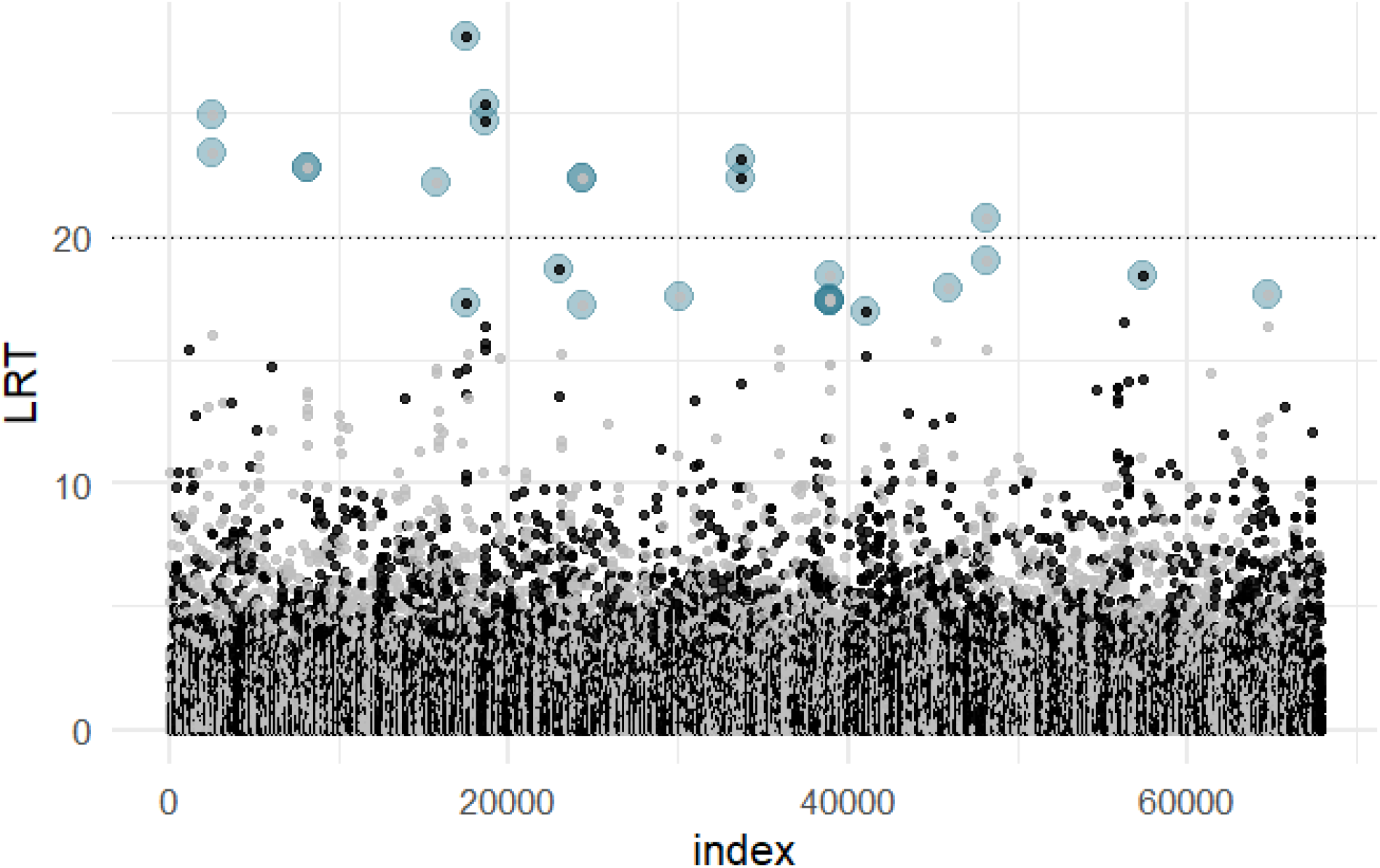
Manhattan of GWAS LRT with SNPs highlighted that are significant outliers. Values above the horizontal line at 20 LRT are significant via both FDR (q-value and Benjamin Hochberg) methods.

### Cochran-Mantel-Haenszel (CMH) test

I found thousands of loci changed significantly across all three treatment replicates in response to selection, with 10,523 in low salinity and 14,322 in high salinity. These represent 0.7% and 1% of the number of loci tested (1.3 million) and 15% and 21% the 68k loci tested in the GWAS, respectively. Of the 39 significant GWAS candidates 2 changed in frequency significantly in low salinity, 4 in high salinity, and 1 in both low and high (Figure 4). Across all significant loci in the CMH test, 42% were unique to high salinity, 37% were shared in both low and high, and 21% were unique to low salinity.

**Figure 4.**
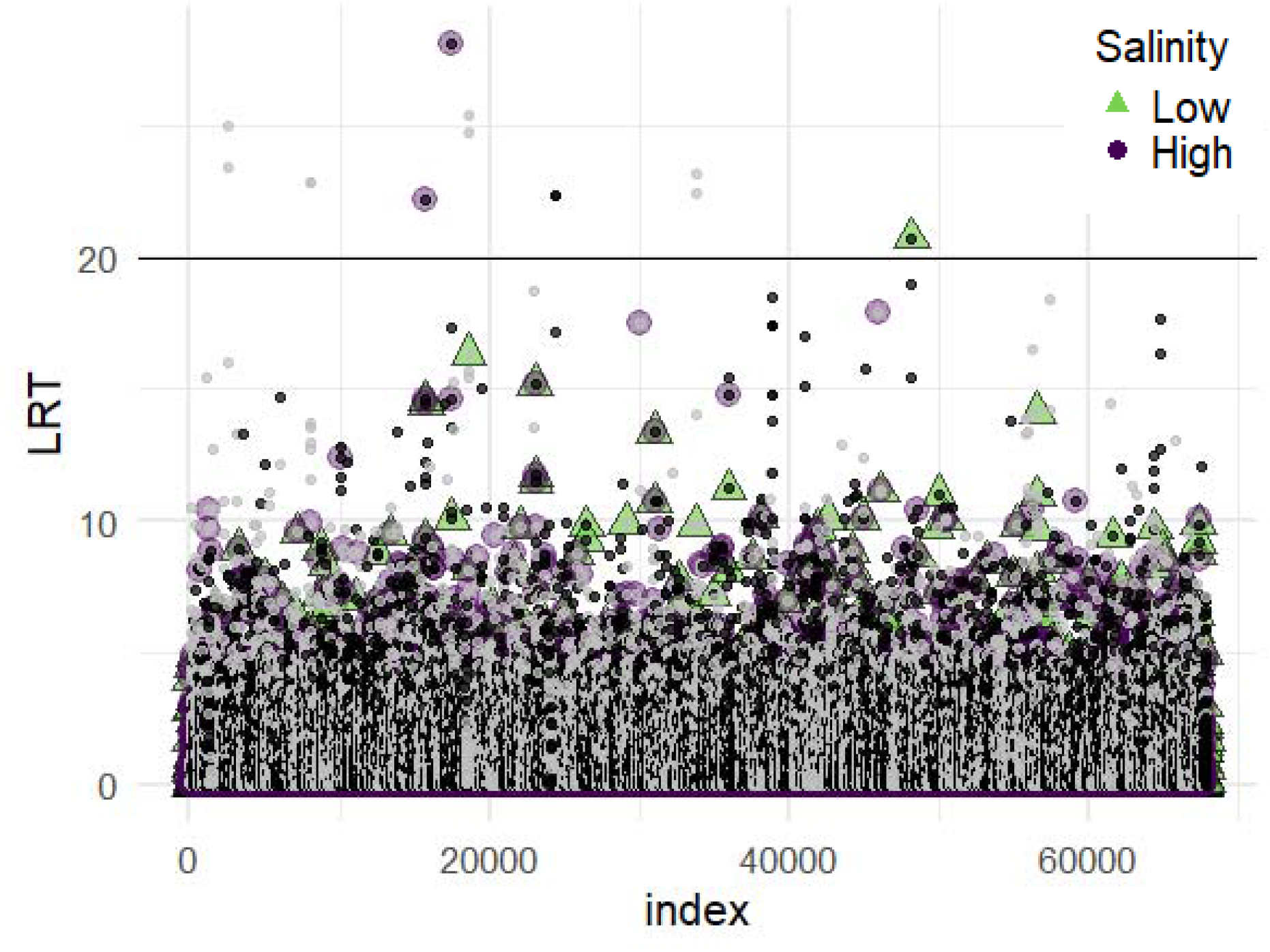
Manhattan of GWAS LRT with SNPs highlighted that are significant outliers in the CMH test of change in allele frequency after selection. Green triangles are the low salinity treatment and purple circles are high salinity.

**Figure 5.**
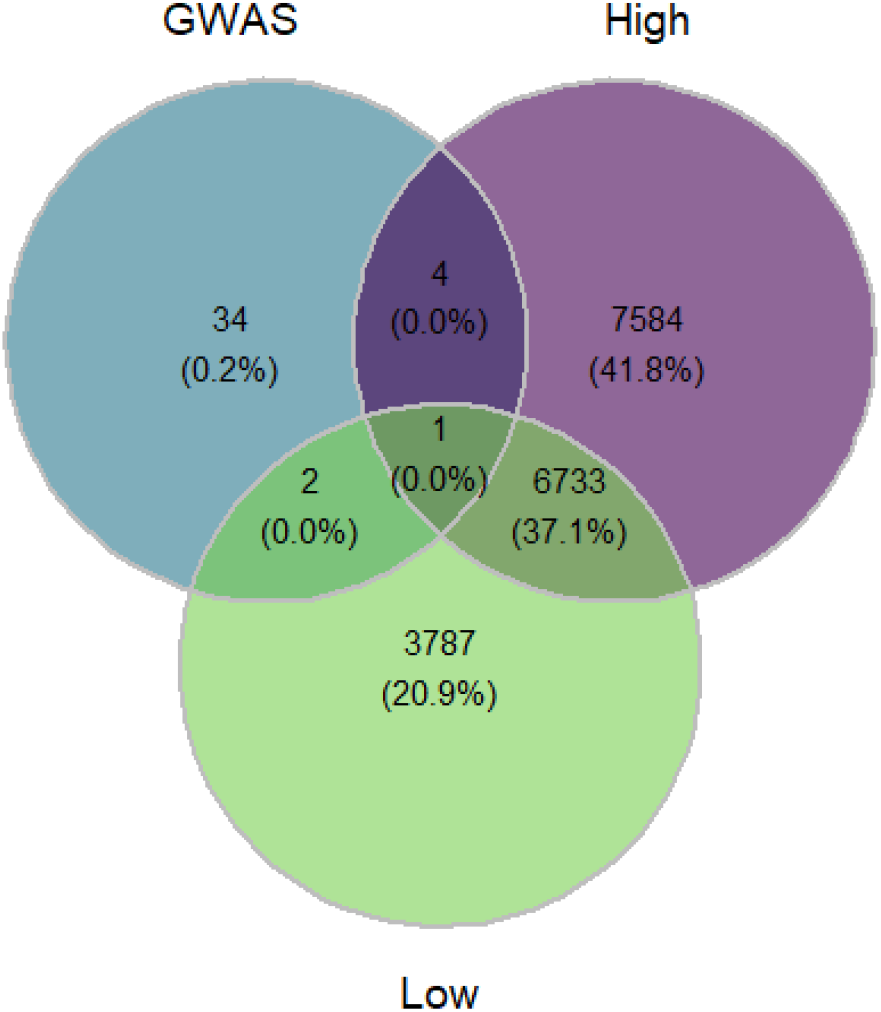
Venn diagram showing relationship of candidate SNPs found in the GWAS, with those found to have significantly changed in frequency after selection in low and high salinity (via the CMH test).

### Reference genome assembly

I extracted high molecular weight DNA that had an average fragment size of 27 kb and which was submitted to the UC Davis Genome Center for library preparation and sequencing. I obtained 793.94 million reads (ca. 120X coverage) for an estimated genome size of 902.38 Mb. Supernova assembly software is optimized to assemble genomes with ∼56X coverage, thus only 336 million reads were used for assembly, as input by the *maxreads* parameter in Supernova. The assembly contig N50 was 24.03 Kb and the N50 scaffold size was 111.64 Kb. In BUSCO against the metazoan database, I found the genome to be 71% complete, 65% complete single-copy, 6% complete duplicated copy, 7% fragmented, and 21% missing.

## Discussion

I found a replicated increase in the proportion of lecithotrophic egg-masses in both low and high salinity and documented heritable variation for offspring-type in the sacoglossan sea slug *Alderia willowi* which exhibits remarkable flexibility for offspring type modulate by environmental conditions. In a genome wide association test, I found 41 loci in 22 genomic contigs that are associated with developmental mode. These results signify a polygenic basis of inheritance for developmental mode in *A. willowi*. A evolve and resequence experiment (E & R) revealed thousands of loci that changed in frequency following selection. Many loci were shared between salinity treatments, and yet many were also unique to the environment in which selection for offspring-type took place. Only a few of the loci found by the GWAS were also found to change significantly in the E & R experiment, suggesting the experiments here may be under-powered to detect more GWAS loci due to small sample sizes, a short number of generations (5) and a strong effect of phenotypic plasticity, which can move the phenotype independent of the genotype.

Evolve and resequence studies enable the use of short term selection experiments on naturally occurring genetic variants to discover the genetic basis of traits under selection, traits that may be correlated with those under selection, and environmental factors that influence trait expression and trait evolution (Long et al. 2015). Furthermore, E & R studies, through the use of replicate populations or selection lines, allow the researcher to determine the relative roles of genetic drift, antagonistic pleiotropy, and positive pleiotropy, as the experimental populations respond to selection (Long et al.

2015, see Gompert and Messina 2016). This approach has revealed interesting levels of variation in how experimental populations respond to selection. For example, in the legume beetle a genetic trade-off, particularly antagonistic pleiotropy, was implicated in resulting in a non-replicated response to host plant switching once the effect of drift was ruled out (Gompert and Messina 2016). Likewise, populations of *Drosophila melanogastor* show a similar phenotypic response to selection for body size but are variable in the number and location of loci which respond to selection (Turner et al. 2011). These studies aid in our understanding of the repeatability of evolution for polygenic traits, with implications on the constraints and genetic trade-offs that may be population or lineage specific.

Developmental mode is a key life-history feature in marine invertebrates which through impacting larval dispersal can influence micro- and macro-evolutionary processes and patterns (i.e, gene flow, demography, local adaptation, speciation and extinction). The sacoglossan sea slug *Alderia willowi* exhibits dramatic variation in life-history, especially during early, critical stages of development, where eggs fall into two discrete size classes, and larval developmental mode varies in concert. I have previously shown that developmental mode in *A. willowi* is influenced by an interaction between genetics and the salinity environment (Caplins, 2021). The results of the selection experiment and GWAS in this paper are consistent with a polygenic basis of inheritance for developmental mode and are consistent among other species of sacoglossan sea slug. In genetic crosses in *Elysia chlorotica* a species which exhibits intraspecific variation in developmental mode as a fixed polymorphism (individuals are not plastic) found in the F1 and F2 generations non-mendelian ratios of adults laying lecithotrophic to planktotrophic egg masses *(West H*.*H, Harrigan J*.*F*., *Pierce S*.*K. 1984)*. A polygenic basis of inheritance in sea slugs is counter to the several large effect genes that determine underlie developmental mode in the poecilogonous annelid *Streblospio benedicti* (Levin, Zhu, and Creed 1991; Zakas and Wares 2012; Zakas et al. 2018). In natural populations of *S. benedicti* maternally determined egg size is associated with specific larval characters, particularly the presence of chaetae that are used for swimming and feeding in the planktotrophic larvae but are absent in the lecithotrophic larvae. In lab crosses these traits can be uncoupled, for example, producing large lecithotrophic larvae with swimming chaetae, and QTL mapping has identified a handful of genetic regions behind egg size and larval phenotype (Zakas et al. 2018). In *S. benedicti* the expression of developmental mode does not appear to be environmentally modulated, and the fitness of alleles that impact zygotic phenotype depend on the maternal background reducing the number of larvae in nature that have an intermediate phenotype (i.e., large lecithotrophic larvae with swimming chaete; Zakas et al. 2018).

---

**Figure S1.**
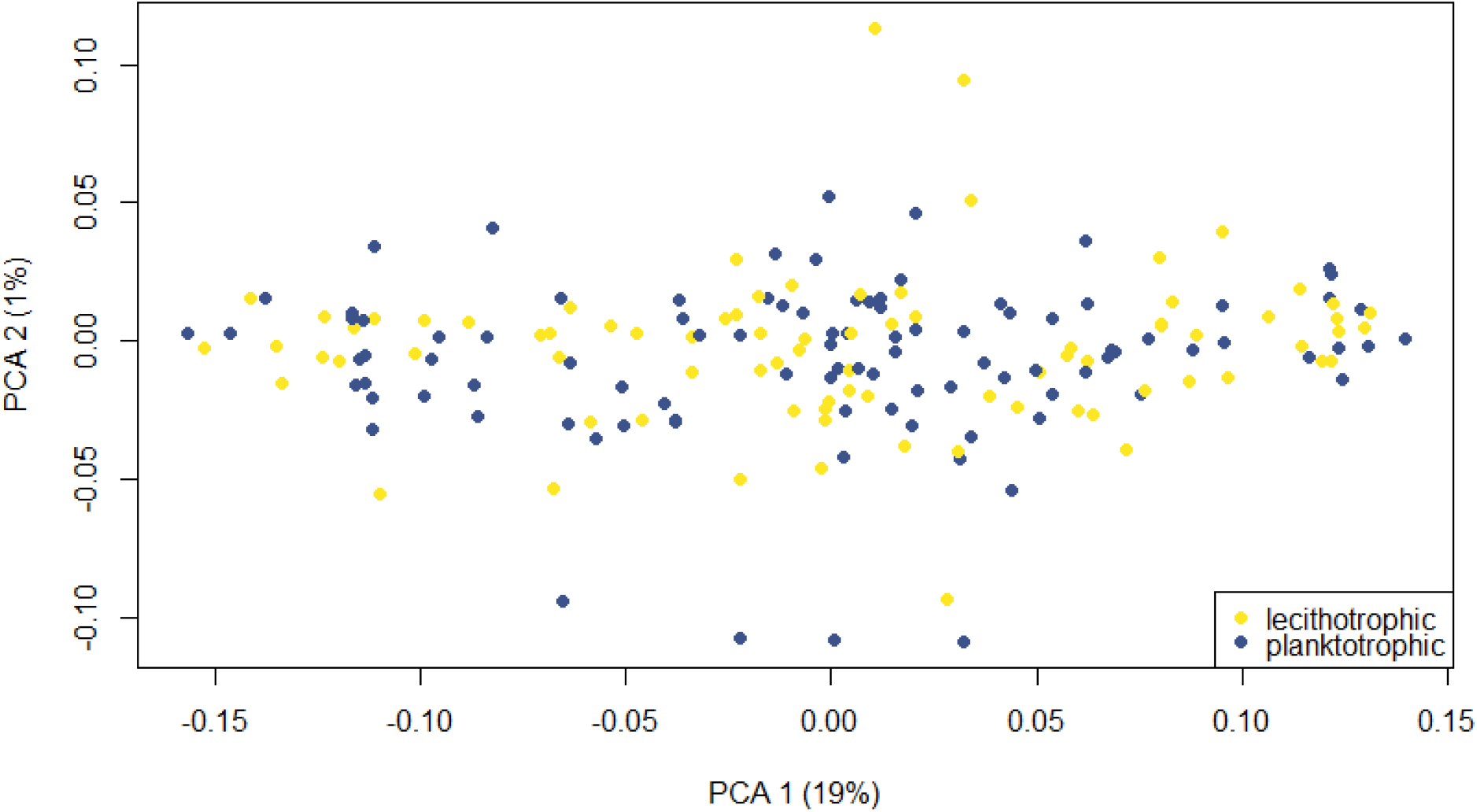
PCA of pre-selection F1s. The lecithotrophic laying slugs are in yellow and the planktotrophic laying slugs are in blue. There is no clear division between the two when considering variation along the subset of the genome for which I have Rad-seq information.

**Figure S2.**
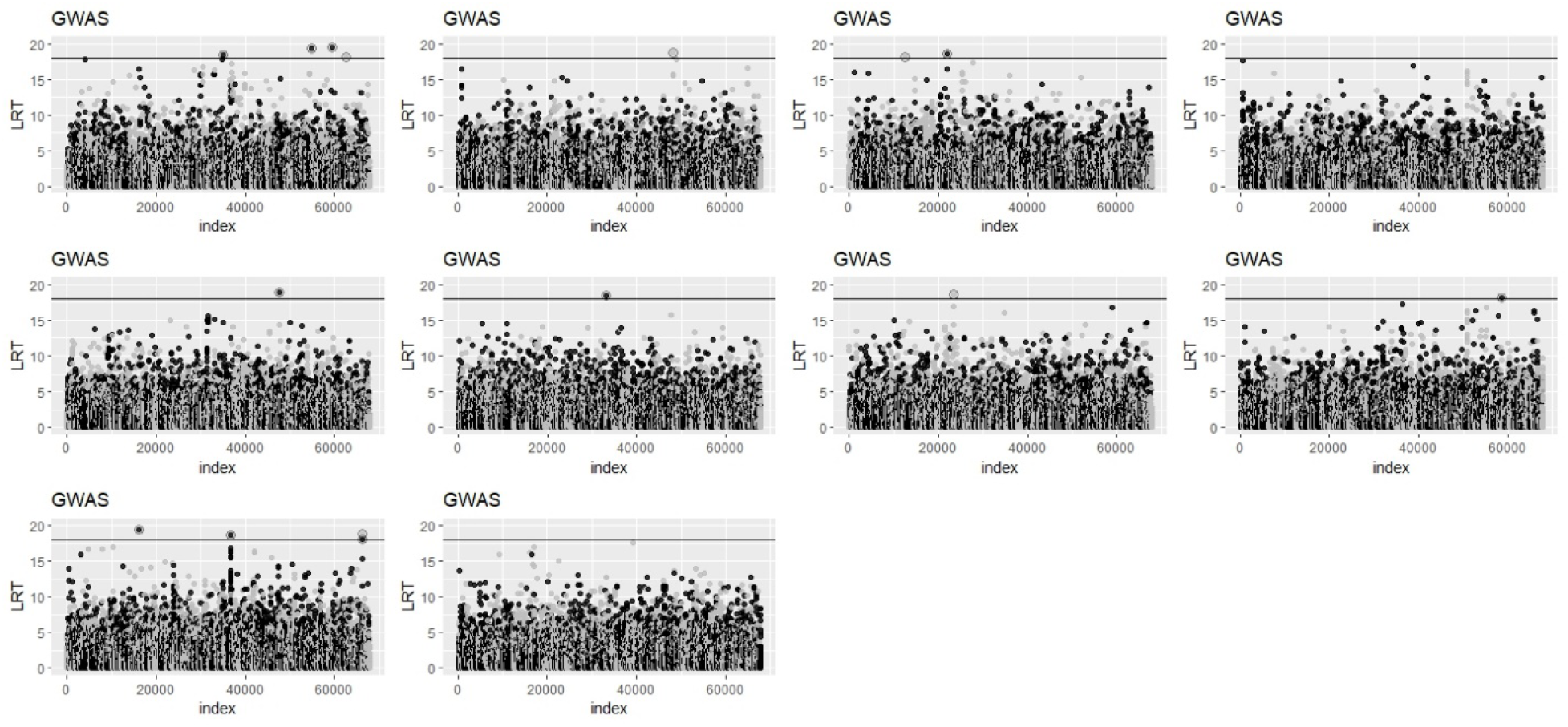
Manhattan plot of likelihood ratio statistic for a genome wide associate test using the genomic data from 187 individuals with a simulated randomly assigned phenotype (repeated 10X). Horizontal line is at LRT =18. Most random tests found 1-2 SNPs associated with the random phenotype and one found 4 with an LRT greater than 18.

